# Mutational scanning of *CRX* classifies clinical variants and reveals biochemical properties of the transcriptional effector domain

**DOI:** 10.1101/2024.03.21.585809

**Authors:** James L. Shepherdson, David M. Granas, Jie Li, Zara Shariff, Stephen P. Plassmeyer, Alex S. Holehouse, Michael A. White, Barak A. Cohen

## Abstract

Cone-Rod Homeobox, encoded by *CRX*, is a transcription factor (TF) essential for the terminal differentiation and maintenance of mammalian photoreceptors. Structurally, CRX comprises an ordered DNA-binding homeodomain and an intrinsically disordered transcriptional effector domain. Although a handful of human variants in *CRX* have been shown to cause several different degenerative retinopathies with varying cone and rod predominance, as with most human disease genes the vast majority of observed *CRX* genetic variants are uncharacterized variants of uncertain significance (VUS). We performed a deep mutational scan (DMS) of nearly all possible single amino acid substitution variants in CRX, using an engineered cell-based transcriptional reporter assay. We measured the ability of each CRX missense variant to transactivate a synthetic fluorescent reporter construct in a pooled fluorescence-activated cell sorting assay and compared the activation strength of each variant to that of wild-type CRX to compute an activity score, identifying thousands of variants with altered transcriptional activity. We calculated a statistical confidence for each activity score derived from multiple independent measurements of each variant marked by unique sequence barcodes, curating a high-confidence list of nearly 2,000 variants with significantly altered transcriptional activity compared to wild-type CRX. We evaluated the performance of the DMS assay as a clinical variant classification tool using gold-standard classified human variants from ClinVar, and determined that activity scores could be used to identify pathogenic variants with high specificity. That this performance could be achieved using a synthetic reporter assay in a foreign cell type, even for a highly cell type-specific TF like CRX, suggests that this approach shows promise for DMS of other TFs that function in cell types that are not easily accessible. Per-position average activity scores closely aligned to a predicted structure of the ordered homeodomain and demonstrated position-specific residue requirements. The intrinsically disordered transcriptional effector domain, by contrast, displayed a qualitatively different pattern of substitution effects, following compositional constraints without specific residue position requirements in the peptide chain. The observed compositional constraints of the effector domain were consistent with the acidic exposure model of transcriptional activation. Together, the results of the *CRX* DMS identify molecular features of the CRX effector domain and demonstrate clinical utility for variant classification.

## Introduction

Cone-Rod Homeobox, encoded by *CRX*, is a photoreceptor-specific transcription factor (TF) essential to both the terminal differentiation of photoreceptors and the maintenance of rod and cone structure and function in adulthood (Swaroop et al. 2010). Over a dozen human sequence variants in *CRX* have been characterized in the pathogenesis of degenerative retinopathies with a wide range of ages of onset and severity, including Retinitis Pigmentosa (RP), a “rod-centric” disease with loss of peripheral vision; Cone-Rod Dystrophy (CoRD), a “cone-centric” disease with loss of central visual field acuity progressing to peripheral vision loss; and Leber Congenital Amaurosis (LCA), an early-onset retinal disease causing severe vision loss and blindness (Freund et al. 1997; Sohocki et al. 1998; Freund et al. 1998; Jacobson et al. 1998; Swaroop et al. 1999; Perrault et al. 2003; Nichols et al. 2010; Huang et al. 2012).

In ClinVar, the NCBI catalog of human gene variants, nearly 150 missense *CRX* variants have been reported, but more than 80% of these are variants of uncertain significance (VUS), with no clinical or functional evidence of their pathogenicity or lack thereof. With the ever-increasing pace of clinical genome sequencing, *CRX* VUS will continue to be identified in patients with degenerative retinopathies, and diagnosis of these patients would benefit from better functional characterization of *CRX* variants.

As a TF, CRX primarily functions by regulating the expression of target photoreceptor genes such as the rod and cone opsins (*RHO*, *OPN1SW*, and *OPN1MW*) (Peng and Chen 2005; Corbo et al. 2010). Pathogenic sequence variants in *CRX* cause profound alterations in the gene expression profiles of photoreceptors that correlate with structural and functional deficits (Tran et al. 2014; Tran and Chen 2014). The E80A variant has been observed in patients with severe dominant CoRD (Freund et al. 1997; Sohocki et al. 1998; Chen et al. 2002), while the nearby K88N variant has been observed to cause dominant LCA (Nichols et al. 2010). The R115Q variant has been reported in patients with Retinitis Pigmentosa (Sohocki et al. 2001), while the R90W variant has been shown to cause LCA when homozygous, but only mild late-onset CoRD when heterozygous (Swaroop et al. 1999; Chen et al. 2002). Pathogenic missense variants have been identified in both the DNA-binding domain and transcriptional effector domain (Figure 1A).

**Figure 1:**
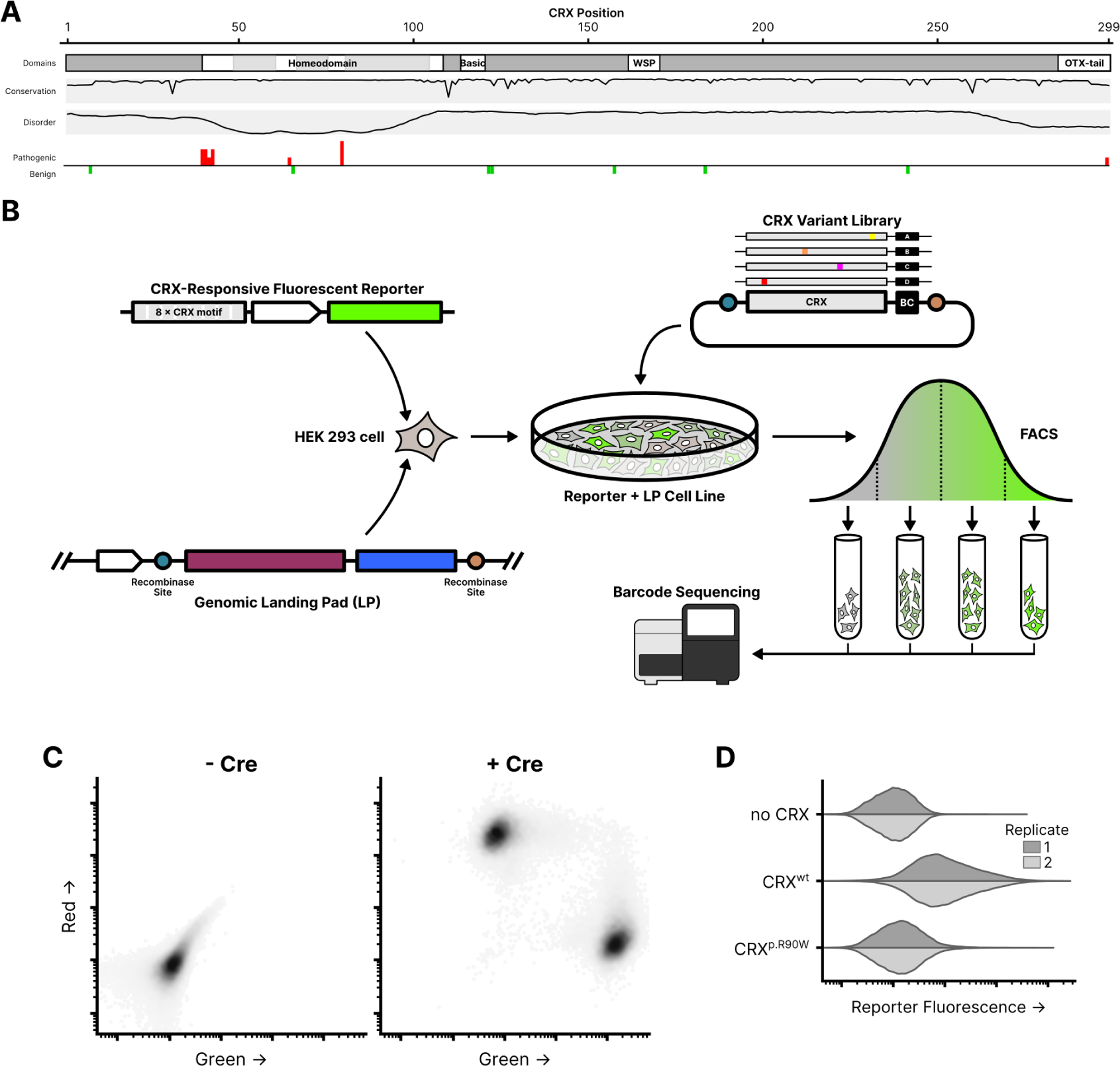
Experimental overview of the CRX deep mutational scan. (**A**) Known CRX domains, sequence conservation, predicted disorder, and reported ClinVar missense variants. Per-residue conservation computed using sequences from the UniProt UniRef50 cluster derived from human CRX. Disorder predicted with Metapredict (Emenecker et al. 2021). Missense pathogenic variants (“Pathogenic” and/or “Likely pathogenic”) and benign variants (“Benign” and/or “Likely Benign”) from ClinVar (accessed 2023-12); height of bar proportional to number of variants at each position (max=3). (**B**) Schematic of the CRX deep mutational scan. A clonal cell line carrying a CRX-responsive fluorescent reporter and genomic landing pad (LP) was generated, and a library of CRX variants was integrated into the LP so that each cell expresses a single variant. Following fluorescence-activated cell sorting (FACS), sequencing was used to determine relative variant barcode abundances in each fluorescence bin, allowing for the calculation of a reporter activity score. (**C**) LP cells integrated with a 1:1 ratio of plasmids carrying mEmerald (green; arbitrary units) or mCherry2 (red; arbitrary units), with or without a plasmid expressing Cre recombinase (60,000 cells plotted per condition, points shaded by density). (**D**) Reporter activation (green fluorescence; arbitrary units) measured in Reporter + LP cells with the indicated CRX variants integrated. Two independent biological replicate experiments per sample (distributions plotted from 40,000 cells).

We applied deep mutational scanning (DMS), which uses libraries of gene variants combined with pooled high-throughput assays, to measure the functional consequences of all missense *CRX* variants in a single experiment. Fundamentally, DMS is the combination of a DNA library of many sequence variants, a method for introducing those variants into a model system in a manner such that the effects of each variant can be individually measured, and a functional assay capable of quantifying the activity of each variant. We developed a cell line carrying a synthetic fluorescent CRX transcriptional reporter and a genomic landing pad (LP) system for controlled single-copy expression of variants. We integrated a plasmid library encoding all single amino acid substitution CRX variants into this cell line so that each individual cell carried a single CRX variant expressed from the LP, sorted cells into bins based on reporter fluorescence, and sequenced variant enrichment across bins to calculate a quantitative activity score for each variant.

We analyzed the DMS variant activity scores in the context of the CRX DNA binding domain structure, the residue composition of the intrinsically disordered transcriptional effector domain, and for the purposes of clinical variant classification. This work provides both a clinical resource for the classification of *CRX* variant effects as well as insight into the rules governing the composition and structure of transcription factors.

## Results

To systematically measure the effects of all variants in *CRX*, we developed a clonal HEK 293-derivative cell line carrying two genomic integrations: a synthetic CRX transcriptional activity reporter, and the aforementioned LP system. The transcriptional reporter contains eight consecutive repeats of a strong consensus CRX binding motif upstream of a CMV-T6 minimal promoter and the fluorescent protein mEmerald (Loew et al. 2010). The LP contains a strong constitutive CMV promoter with asymmetric lox sites downstream (Figure 1B). Upon the addition of a plasmid with matching lox sites and Cre recombinase, the LP can undergo recombinase-mediated cassette exchange (RMCE), allowing for the precise insertion of a CRX variant at the defined genomic LP locus, under the control of the CMV promoter. HEK 293 cells were used due to their experimental tractability, and a history of prior successful reporter assays for CRX (Chen et al. 2002; Peng et al. 2005; Tran et al. 2014). The endogenous *CRX* locus is repressed and unexpressed in HEK 293 cells.

The use of a genomic LP achieves precise expression of the integrated gene cassette, allowing for the expression of a single CRX variant in each cell. To validate the LP cell line, we co-transfected a 1:1 mixture of two plasmids, carrying either the mEmerald or mCherry2 fluorescent proteins with flanking compatible lox sites. Without the addition of a plasmid expressing Cre recombinase, the fluorescent proteins were not expressed, showing that RMCE is required for protein production (as the plasmids themselves lack a promoter). With the addition of Cre recombinase, cells could be observed with tightly controlled green or red fluorescence but not the expression of both colors, demonstrating that constructs could be successfully expressed upon RMCE and that the cell line only harbored a single LP (Figure 1C).

To demonstrate the sensitivity of the synthetic CRX transcriptional reporter, we separately integrated three different constructs into the genomic LP: a transcriptionally-inactive control construct (the miRFP670 fluorescent protein), wild-type CRX, or the known hypomorphic p.R90W CRX variant (Figure 1D). HEK 293 cells do not natively express their endogenous copy of *CRX*, minimizing background protein expression that could interfere with assay measurements (Figure S1). As expected, wild-type CRX alone was capable of activating the reporter construct and driving expression of the fluorescent reporter protein, while the p.R90W variant protein failed to activate the reporter above background fluorescence as compared to the inactive control.

### A cell-based CRX reporter system quantitatively measures variant activity

We cloned a library of all possible single amino acid substitution *CRX* variants, and associated each variant with a unique random barcode sequence using PacBio long-read sequencing (Figure S2A). This library was integrated into the clonal cell line carrying the CRX reporter and genomic LP by RMCE. The fluorescence of each cell is proportional to the level of reporter activation by the CRX variant expressed in that cell. We sorted the cells into four bins based on fluorescence, extracted genomic DNA, and sequenced variant barcodes. The relative abundance of each barcode in each bin, and thus the relative abundance of each variant in each bin, were used to calculate per-variant activity scores (Figure 1B, Figure S2B-C). After filtering and quality control for low representation or read abundance, we were able to calculate activity scores for 5,285 variants out of 5,662 designed variants.

The computed variant activity scores for each variant show patterns of sensitivity to substitutions that correspond with different structural domains of CRX (Figure 2). Residues in the homeodomain are particularly sensitive to substitution, as seen in the average variant effect track. In the central region of CRX (residues ∼95–150), apart from the basic domain, residues are largely insensitive to substitution. In the remainder of the disordered regions (including at the N-terminus), substitutions for specific classes of amino acids (such as negatively charged residues or aromatics) show specific increases or decreases in activity. In the C-terminus, substitutions of a number of residues tend to substantially affect CRX transcriptional activity, with substitutions of specific residues systematically increasing or decreasing activity. Below, we discuss each of these trends in more detail.

**Figure 2:**
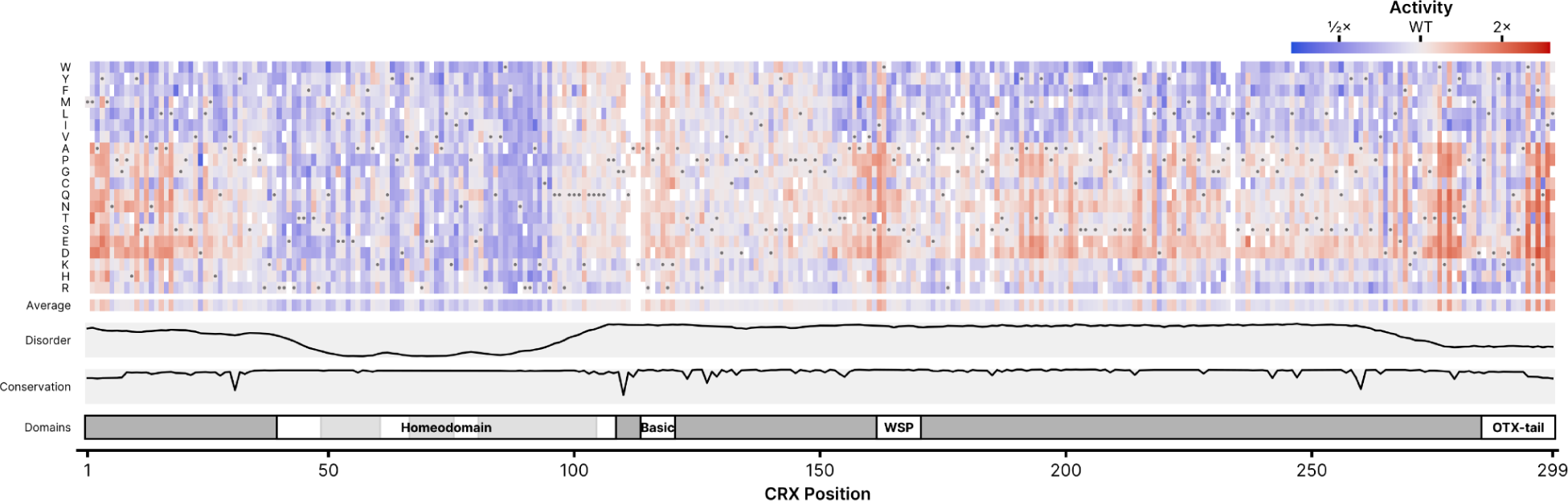
DMS activity scores for all measured single amino acid CRX substitutions. Activity scores were normalized to wild-type CRX; the wild-type amino acid at each position is indicated by the gray circle in each column. The average row shows the mean activity score for all substitutions at each position. Disorder, domains, and conservation are shown as in Figure 1. Empty boxes indicate variants not measured in the DMS assay, due to drop-out during the library cloning or variant measurement steps.

### Activity scores can be used to classify known pathogenic CRX variants with high specificity

Although only nineteen missense *CRX* variants have been unambiguously classified in ClinVar (Figure 1A), we can use the activity scores for these variants to benchmark the classification performance of the DMS assay. The DMS activity scores for ClinVar variants are shown in Figure 3A. Some variants with “Conflicting” classifications in ClinVar can be confidently manually-curated into the “Benign” or “Pathogenic” categories by literature review and are indicated in the figure. Although the scores for the benign variants generally cluster around wild-type, the scores for the pathogenic variants are more varied. To assign a statistical confidence to each variant, we took advantage of the design of the DMS assay—specifically, that each variant is marked by multiple sequence barcodes. The distributions of barcode-level activity scores for several selected variants are shown in Figure 3B. We leveraged the fact that in the *CRX* variant library, wild-type *CRX* is barcoded thousands of times. We used a two-sample Kolmogorov–Smirnov (K-S) test to compare the shapes of each variant’s barcode-level activity distribution to that of wild-type CRX. Combining the DMS activity scores with the FDR-corrected K-S p-value, we were able to set cutoffs for variants with significantly low or high activity (Figure 3C).

**Figure 3:**
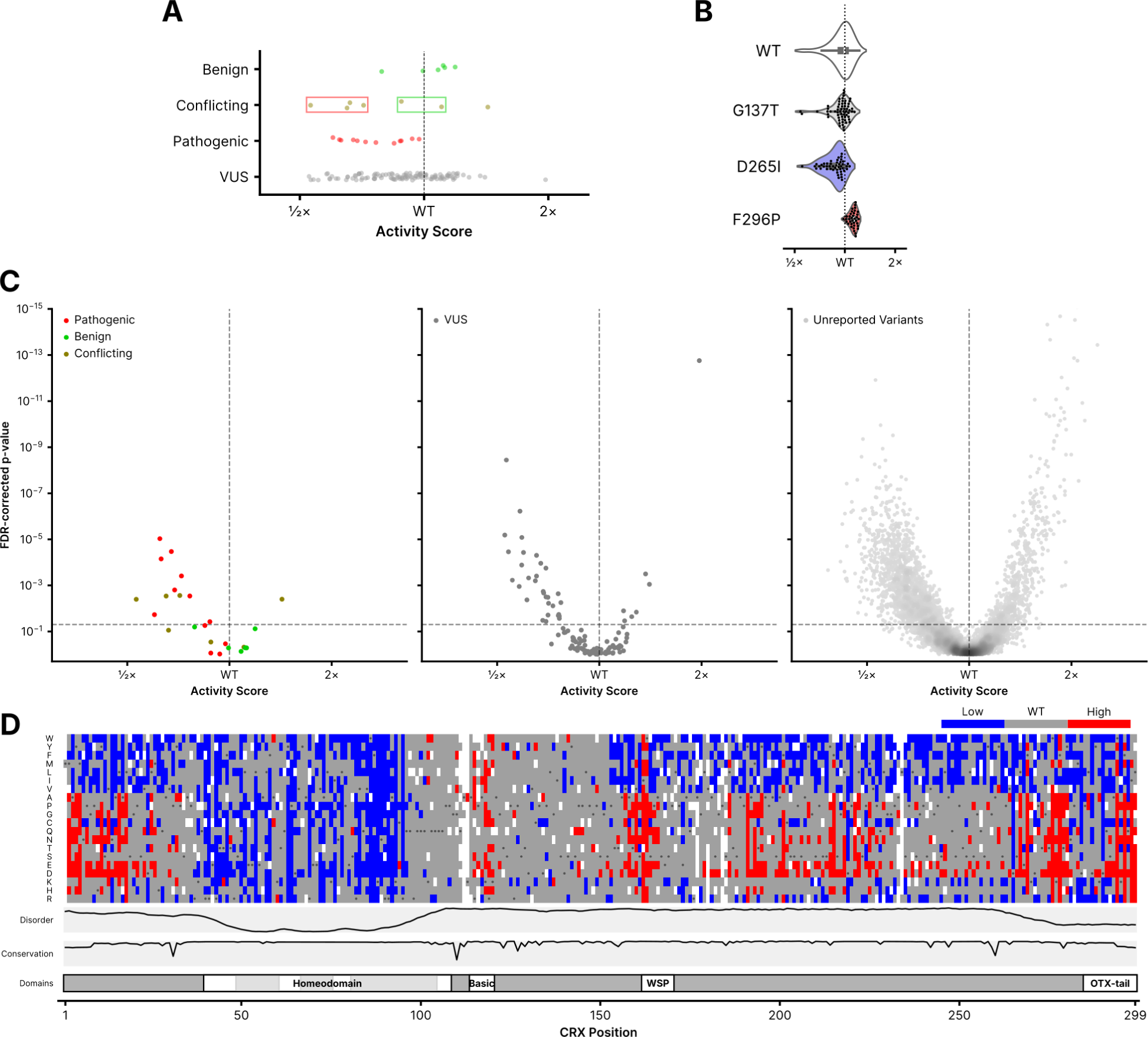
Classification of CRX variants. (**A**) DMS activity scores for variants reported to ClinVar in each of the indicated classes (Pathogenic includes “Pathogenic” and/or “Likely pathogenic”; Benign includes “Benign” and/or “Likely Benign”). For variants in the Conflicting Interpretations class, reanalysis of the boxed variants support re-classification into the groups indicated by box color. (**B**) Barcode-level activity measurements for wild-type CRX and the indicated representative wild-type-like (p.G137T), low-activity (p.D265I), and high-activity (p.F296P) variants. Each black dot represents a unique barcoded construct; not shown for wild-type CRX due to it being barcoded thousands of times. (**C**) Volcano plots showing classifications for the indicated ClinVar variants (left and middle panels) or all other variants not yet reported in ClinVar (right panel). FDR-corrected p-values were computed from a two-sample Kolmogorov–Smirnov test comparing each variant’s barcode-level measurements to those of wild-type CRX, as visualized in (B). The horizontal line corresponds to an FDR-corrected p-value of 0.05; the vertical line corresponds to a normalized activity score of 1 (wild-type). For visualization purposes, the y-axis is clipped to 10^-15^; 13 variants are hidden with activity scores greater than wildtype and p-values up to 10^-26^. (**D**) Quantized DMS activity scores for each variant, coloring low- and high-activity variants using the significance cutoffs shown as dotted lines in (C). Disorder, domains, and conservation are shown as in Figure 1.

Using these criteria, we see that the DMS assay is a high-specificity pathogenicity classifier: all eight benign (or “Conflicting” reclassified to benign) variants have DMS activity scores that are not significantly different from wild-type CRX. For the 16 pathogenic (or “Conflicting” reclassified to pathogenic) variants, 11 show significantly-reduced activity scores. Using the significance criteria, we generated a list of high-suspicion activity-altering variants (Figure 3D). When combined with other clinical data sources, this list of variants identifies substitutions with the highest predicted impact on CRX activity, and thus greatest likelihood for pathogenicity. Using the OddsPath framework (Tavtigian et al. 2018) for benchmarking functional assay data for variant classification, this corresponds to an OddsPath score of 5.5, meeting the criterion for the PS3_Moderate evidence level in the ACMG variant classification framework (Brnich et al. 2020). Because of the small number of reported CRX variants in ClinVar, PS3_Moderate is the maximum achievable evidence level even for an assay with perfect variant discrimination.

### Variants can affect protein abundance, but abundance alone does not explain changes in activity

We selected a group of low- and high-activity variants to individually validate in one-at-a-time integration experiments (Figure S3A). For these variants, we also performed western blots to assess their effects on CRX protein abundance. For each variant, we assessed blot intensity relative to a β-tubulin control for each sample. In general, low-activity variants tended to show reduced abundance compared to wild-type CRX while high-activity variants showed increased abundance (Figure S3B). However, abundance alone was not enough to explain the activity of all variants, as some low-activity and high-activity variants had wild-type-like abundance. This analysis is also limited by the fact that western blot quantification can be a noisy measurement of protein abundance, but suggests that protein abundance effects may explain a portion of the observed changes in variant activity.

### Variant activity scores in the homeodomain closely align with a predicted structural model

To benchmark our assay results and evaluate how closely they reflect known physical constraints of CRX-DNA binding, we mapped the average per-residue DMS activity scores onto a structure of the CRX homeodomain in complex with DNA (Figure 4A). Because an experimentally-determined structure of CRX is not available, we aligned the AlphaFold2-predicted CRX structure to crystal structures of other homeodomains in the paired family, which is highly structurally conserved. Strikingly, DMS activity scores are highly concordant with the structural model. Amino acid residues in the major groove are particularly sensitive to substitution, even when compared to residues in the same α-helix that do not lie in the major groove. Furthermore, residues on the DNA-facing sides of the two α-helices not in the major groove are more sensitive to substitution than residues on the outward-facing sides of the same α-helices.

**Figure 4:**
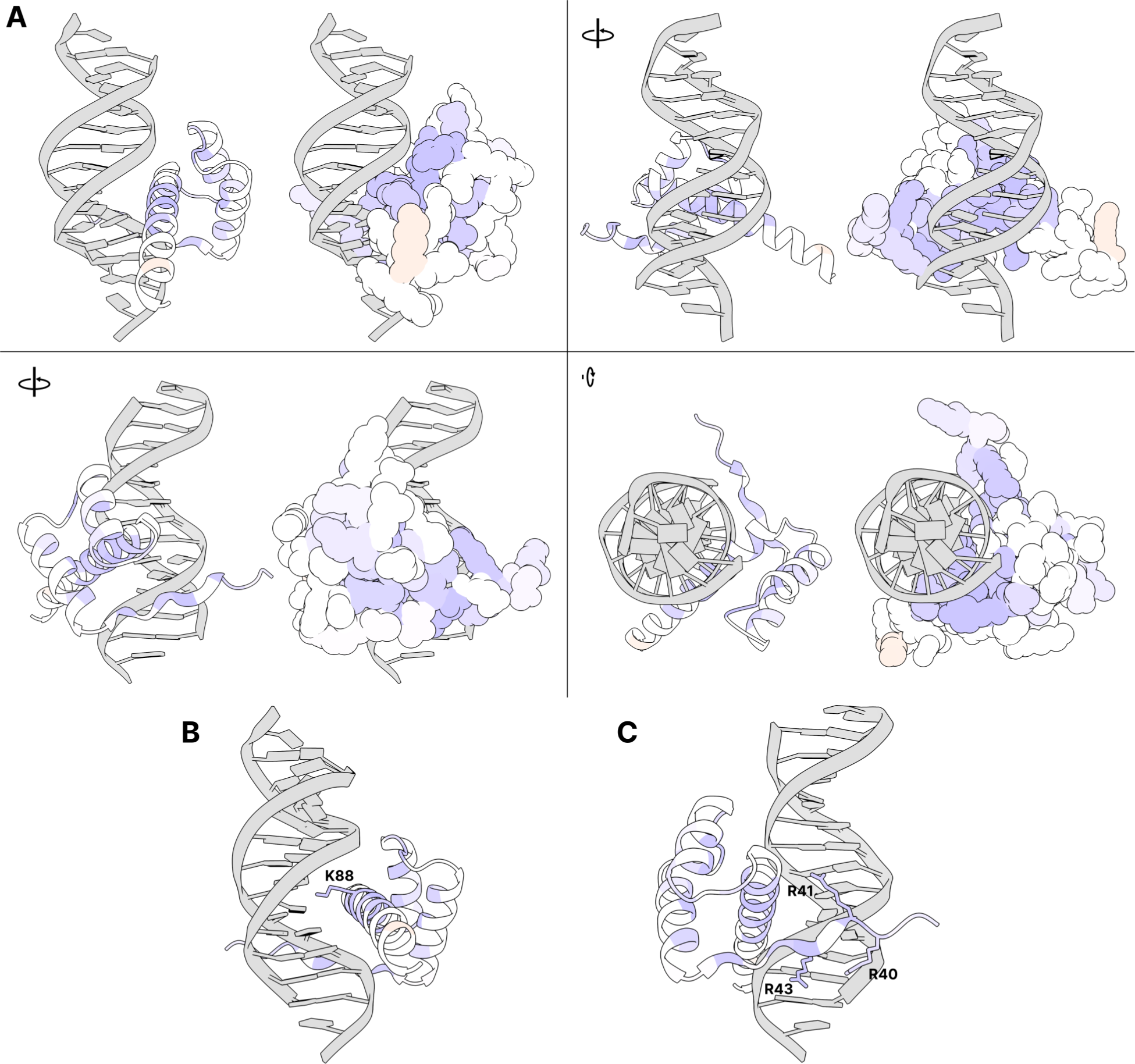
Average DMS activity scores superimposed on a predicted structure of the CRX homeodomain in complex with DNA. (**A**) Various views of residues 38–104 of an AlphaFold predicted structure of CRX aligned to a crystal structure of *Drosophila* paired in complex with DNA (PDB 1FJL) (Wilson et al. 1995). For each view, a cartoon ribbon model is shown on the left and a space-filling atomic model is on the right. (**B**) Close-up of residues in the major groove with K88 highlighted (side chain shown). (**C**) Close-up of minor groove-contacting residues with arginine residues highlighted (side chains shown). In all panels, residues are colored by average DMS activity score, as shown in the “Average” track in Figure 2.

CRX is a K50-type homeodomain transcription factor, which refers to the lysine (K) residue at the 50th position of the homeodomain sequence which is crucial for determining DNA binding specificity (Noyes et al. 2008). In CRX, residue 88 is the K50 position, and substitutions at this position do indeed have particularly large effects on activity (Figure 4B, Figure 2). It is also known that arginine residues at the N-terminus of the homeodomain interact with the minor groove to further control motif specificity (Noyes et al. 2008). These residues are particularly sensitive to substitution as predicted, even when compared to immediately adjacent non-DNA-contacting residues (Figure 4C). Taken together, the strong concordance of the DMS activity scores with the structural model in general and at known crucial residues provides support that the DMS assay is accurately measuring the ability of CRX variants to bind and activate the transcriptional reporter, and provides mechanistic and structural insight into substitution sensitive and non-sensitive amino acid residues. Furthermore, these results lend confidence to the biological relevance of the synthetic HEK 293-based transcriptional reporter used in these experiments.

### Patterns of substitution activity in the transcriptional effector domain support an acidic exposure model

In addition to its use in classifying clinical *CRX* variants, the DMS assay reveals molecular features of the disordered CRX transcriptional effector domain. Throughout the effector domain, substitutions that add negative charge (aspartic acid or glutamic acid) tend to increase activity, while substitutions that remove existing negatively charged residues (particularly D24, D219, D265, D271, D284, D287, and D290) tend to reduce activity. Conversely, substitutions that add aromatic residues (phenylalanine, tryptophan, and tyrosine) or aliphatic hydrophobic residues with larger side chains (methionine, leucine, isoleucine, and valine) tend to decrease activity, while substitutions that remove existing residues in these classes tend to increase activity, especially when replaced with polar uncharged or negatively charged residues (Figure 2).

Further underscoring the composition-sensitivity of the effector domain, the effects of residue substitutions can be used to cluster amino acids by the chemical properties of their side chains. We performed unsupervised hierarchical clustering by substituted residue on positions in the disordered region of the transcriptional effector domain (residues 2–38 and 153–264, Figure 5A). Substituted amino acids largely cluster by their biochemical properties, supporting the idea that the disordered transcriptional effector domain is primarily sensitive to overall amino acid composition, rather than the identity of particular residues at certain positions.

**Figure 5:**
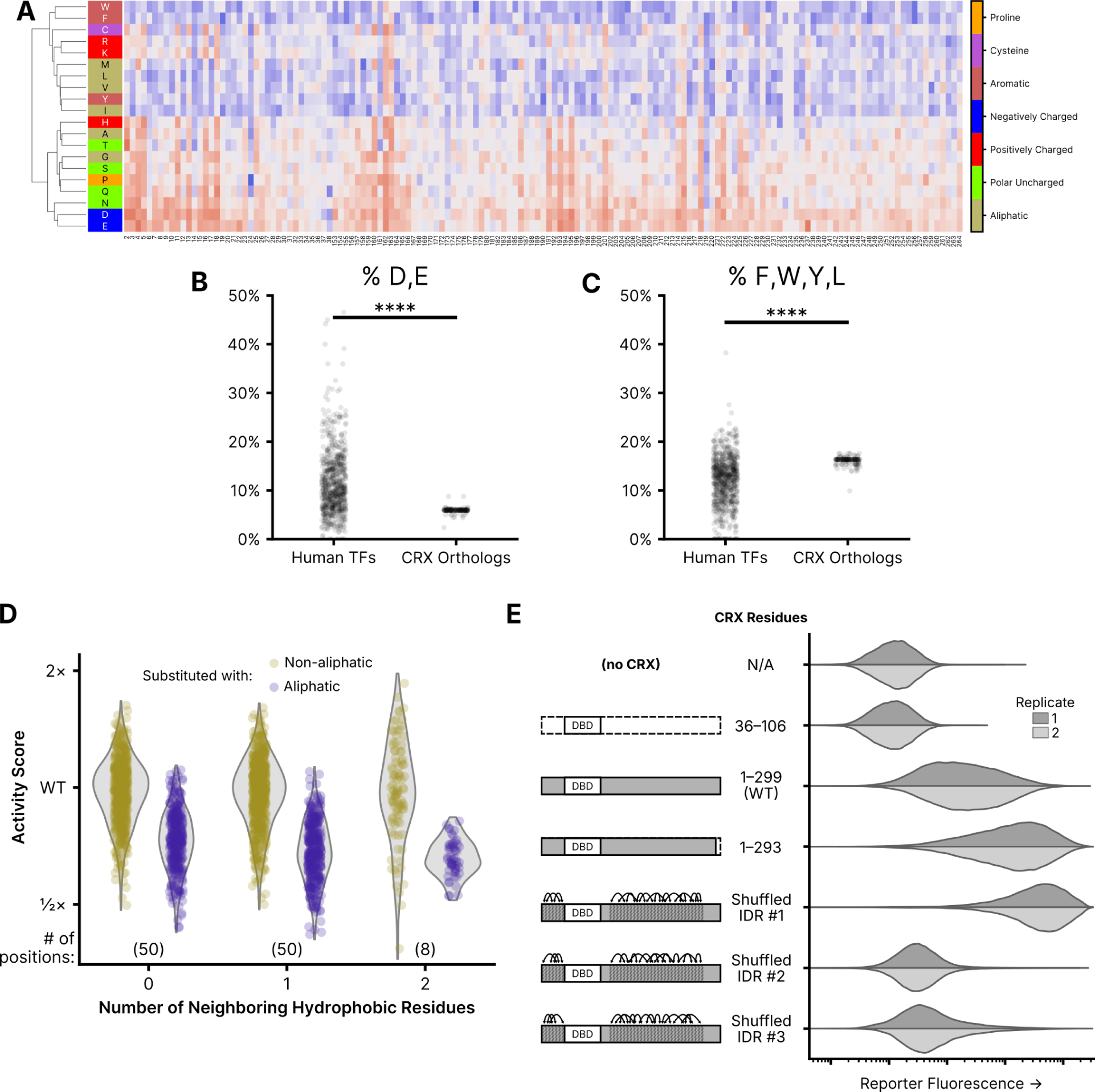
Residue class preferences in the CRX transcriptional effector domain. (**A**) Unsupervised hierarchical clustering (UPGMA method) of per-position activity scores for residues in the disordered region of the transcriptional effector domain (residues 2–38 and 153–264). Residues colored by class. Substitution activity scores colored as in Figure 2. (**B**) Abundance of aspartic acid (D) and glutamic acid (E) residues in disordered regions of all human TFs and CRX orthologs. (**C**) Abundance of phenylalanine (F), tryptophan (W), tyrosine (Y) and leucine (L) residues in disordered regions of all human TFs and CRX orthologs. For (B) and (C), significance tested by two-sided Mann-Whitney U-test; p < 1e-39. (**D**) Comparison of the effects of substituting non-hydrophobic positions in wild-type CRX with hydrophobic (F, W, Y, M, I, L, or V) or non-hydrophobic amino acids, separated by the number of neighboring hydrophobic residues. Analysis limited to positions in the disordered transcriptional effector domain; the number of positions in each neighbor group is shown in parentheses. (**E**) Reporter activation (green fluorescence; arbitrary units) measured in Reporter + LP cells with the indicated CRX variants integrated. Two independent biological replicate experiments per sample (distributions plotted from 50,000 cells).

The balance of effects between substitutions affecting negatively charged and aromatic residues in the transcriptional effector domain is notable in light of the acidic exposure model of transcriptional activation domains (Staller et al. 2022). The acidic exposure model proposes that strong transcriptional activators are facilitated by a balance of acidic and aromatic residues, in which acidic residues permit the solubilization of hydrophobic motifs. Even amongst transcription factor effector domains, the CRX transcriptional effector domain is particularly low in negative charge and high in aromatic residue content (Figure 5B-C). This is consistent with the observed substitution effects of residues in these classes: substitutions of negatively charged residues to amino acids with uncharged or positively charged side chains uniformly decrease CRX transcriptional activity while substitutions of uncharged or positively charged residues to negatively charged amino acids increase CRX transcriptional activity, and the reverse is true for aromatic residues.

To examine the role of local sequence context in determining the functional effect of hydrophobic substitutions, we compared the activity scores of hydrophobic substitutions (hydrophobic residues defined as F, W, Y, M, L, I, or V) to all other substitutions at non-hydrophobic residues in the transcriptional effector domains (residues 2-38 and 153-264), stratifying positions according to the number of immediately adjacent hydrophobic residues (Figure 5D). We fit an interaction model between the class of substituted residue and the number of neighboring hydrophobic residues, finding that the deleterious effect of hydrophobic substitutions increased with the local density of hydrophobic residues (likelihood ratio test p = 2.37×10^-4^). Though subtle, the context dependence of hydrophobic substitution effects indicates that effector domain function is sensitive to residue spacing as well as composition.

To further test whether the CRX transcriptional effector domain exhibits any residue position requirements or is primarily sensitive to overall amino acid composition, we tested three CRX variants in which we shuffled the position of the amino acids in the disordered region. To shuffle the amino acids, we broke the disordered regions into five residue windows and randomized the order of the residues within each window. This ensures global sequence chemistry is preserved while local sequence order is altered. Two of the three shuffled variants showed reduced transcriptional activity compared to wild-type CRX, but the third showed the strongest reporter activation of any CRX variant tested (Figure 5E). From this result, it is clear that the effector domain is not entirely position-insensitive, as three arbitrary amino acid rearrangements demonstrated substantially different effects on transcriptional activity. Nevertheless, there is clearly a qualitative difference in position sensitivity between the ordered and disordered regions of CRX.

We also tested a variant truncating the six most C-terminal residues of the OTX-tail motif. Substitutions of any of these amino acids (WKFQIL), which include several aromatic and leucine residues, largely increased transcriptional activity in the DMS assay, particularly when replaced with aliphatic or negatively charged side chains. Consistently, the 1–293 truncation variant activates the CRX transcriptional reporter more strongly than wild-type CRX, although not as strongly as the strongest shuffled variant. This result suggests that these residues of the OTX-tail motif may play a repressive role in the CRX transcriptional effector domain.

## Discussion

We conducted a systematic characterization of variant effects in the photoreceptor transcription factor CRX. Using a synthetic fluorescent reporter, we measured the activity of nearly all possible missense variants in *CRX*, and assigned each a functional activity score. Substitutions showed unique patterns of variant activity in different functional domains of CRX. Residues in the DNA-binding homeodomain were particularly sensitive to any substitution, and per-residue average activity scores closely aligned to a predicted structure of the CRX homeodomain in complex with DNA. In the intrinsically disordered transcriptional effector domain, variant effects were primarily dictated by biochemical amino acid class rather than position, and substitution effects were consistent with the acidic exposure model of transcriptional activation domains.

From the observed patterns of variant effect, CRX can be broken into three main regions. The DNA binding domain, roughly spanning residues 40 to 95, is highly sensitive to substitution, and many substitutions result in reduced reporter activity, particularly for DNA-contacting or folding-essential residues. The central intrinsically disordered region of CRX, by contrast, spanning residues 95 to 155, is largely insensitive to substitution, with the exception of some residues in the basic domain that tend to result in increased activity upon substitution. Lastly, the C- and N-terminal disordered regions show strong sensitivity to amino acid class, with both increases and decreases in activity depending on substitution. While disordered regions lack a fixed 3D structure and are therefore often assumed to be relatively insensitive to mutations, our work clearly reveals chemically-interpretable patterns in mutational sensitivity.

As discussed, the acidic exposure model of activation domains suggests that strong transcriptional activators require a balance of acidic residues with aromatic and leucine residues (Staller et al. 2022). Compared to other transcription factor disordered domains, CRX has a high proportion of aromatic and leucine residues in the disordered transcriptional effector domain and a low proportion of acidic residues (Figure 5B–C). This may explain why substitutions increasing negative charge or decreasing aromaticity result in increased activation—the CRX transcriptional effector may have a residue composition that is not the same as that of an “ideal” strong activator. We also observed that the local density of hydrophobic residues in the transcriptional effector domain affected variant activity, with substitutions of hydrophobic amino acids in sites with a higher number of neighboring hydrophobic residues tending to show stronger reductions in transcriptional activity. This effect appears consistent with functional constraints on the relative spacing of aliphatic and aromatic residues observed in other hydrophobic-enriched IDRs (Jonas et al. 2023, Martin et al. 2020, Holehouse et al. 2021), indicating that the introduction of hydrophobic patches in the CRX transcriptional effector domain may impair solubility or accessibility of residues required for activation. CRX and other homeodomain transcription factors have been shown to be capable of acting as both activators and repressors at different target sequences (White et al. 2016; Shepherdson et al. 2024; DelRosso et al. 2023). Further characterization is required, but balancing activation and repression in the transcriptional effector domain may necessitate the “intermediate” activation strength residue composition observed for CRX. If so, the pathogenicity of high-activity variants could be further compounded by impaired repression at particular target cis-regulatory elements.

The CRX transcriptional effector domain also has a high proportion of proline and serine residues. If any of these serine residues were to undergo phosphorylation, they would acquire negative charge that could potentially contribute to the residue balance requirements of the acidic exposure model. Proline-directed protein kinases, including members of the MAPK family, have been shown to target NRL, a key transcription factor expressed in rods and a close partner of CRX (Swain et al. 2007). Furthermore, other protein kinases, including PKC family members, have been reported to be essential for rod cell development in mice (Pinzon-Guzman et al. 2011). Whether these or other protein kinase enzymes interact with CRX and what the impact may be of any possible serine phosphorylation on the CRX transcriptional effector domain will require further study.

The high-specificity classification we observed for clinical variants, with a low false positive rate but moderate false negative rate, is reasonable given the design of the reporter assay. In the genome, CRX binds numerous cis-regulatory elements with varied and non-consensus binding motif sequences (Corbo et al. 2010). The synthetic CRX transcriptional reporter used in this study, which contains only high-affinity consensus motif sequences, may be insensitive to subtle variants that alter DNA binding at non-ideal motifs. Furthermore, it is completely insensitive to variants that cause a gain of novel binding motifs. For instance, the p.E80A CRX variant, a known pathogenic variant in humans, displays wild-type-like activity in the DMS assay. It has recently been shown that while the p.E80A CRX variant prefers the same consensus motif as wild-type CRX, it is more tolerant of base mismatches in the motif, an effect to which the DMS reporter may not be sensitive (Zheng et al. 2023).

Furthermore, the LP + reporter cell line is a synthetic transcriptional environment, lacking many of the partner transcription factors typically expressed in photoreceptors. In the retina, CRX is known to co-bind at cis-regulatory elements with a number of TFs, including NRL, OTX2, RAX, RORB and others (Friedman et al. 2021). How the presence or absence of these factors influences the functional consequences of *CRX* variants remains to be seen. However, the fact that ectopic expression of CRX variants and the addition of a single transcriptional reporter construct to a non-photoreceptor cell type is enough to produce data so consistent with existing structural data, variant classifications, and activation domain models is a testament to the power of the reductionist approach used in this DMS assay.

The approach used in this study, with a synthetic transcriptional reporter and non-native cell type, may be broadly applicable to other TFs with expression limited to specific cell types that are difficult to obtain or culture. Compared to the growth and survival assays that are more typical of deep mutational scans in mammalian cells (Maes et al. 2023), transcriptional reporters have the benefit of being a direct readout of TF protein activity. This may help to increase their sensitivity to the effects of protein variants, even in a non-physiologic context.

That a CRX variant with shuffled residues in the intrinsically disordered transcriptional effector domain was capable of activating the reporter more strongly than wild-type CRX further supports that *CRX* did not evolve to encode the strongest possible activation domain, consistent with the discussion above of a potential balance of activation and repression. Furthermore, the varied effects on transcriptional activity observed across the three shuffled variants suggest that the precise residue order has an effect on activation strength, even if specific positions are generally less individually sensitive to substitution than those in the ordered and folded homeodomain. This conclusion is in line with other analyses of other DNA-binding proteins (Langstein-Skora et al. 2022; Sanborn et al. 2021; Jonas et al. 2023; Mindel et al. 2023).

The variant activity scores in the intrinsically disordered transcriptional effector domain are particularly relevant in light of recent computational variant effect predictors trained on protein sequence homology and structural data. AlphaMissense (Cheng et al. 2023), ESM1b (Brandes et al. 2023), and EVE (Frazer et al. 2021) use machine learning approaches to predict variant pathogenicity from, primarily, evolutionary protein sequence conservation data. This is of particular relevance to proteins with intrinsically disordered regions, however, because disordered proteins are not subject to the same evolutionary sequence constraints as structured proteins (Holehouse and Kragelund 2024). Visualization of variant classifications derived from the DMS activity scores compared to classifications from AlphaMissense, ESM1b, and EVE show concordance in the structured homeodomain, but there is less agreement in the disordered transcriptional effector domain (Supplementary Figure S4). Future computational variant effect predictors would likely benefit from the incorporation of functional mutational scanning data, particularly for proteins with intrinsically disordered regions like CRX.

The limitations of this work primarily relate to the intrinsic limitations of reporter assays in non-primary cell types. As discussed, the synthetic transcriptional reporter developed in this study cannot capture the full breadth of CRX-bound cis-regulatory elements in the genome. Furthermore, while extremely tractable, the HEK 293-derived cell line used for these experiments does not reflect the unique transcriptional environment of photoreceptors. For instance, the substitution-tolerant central region of CRX could conceivably be necessary for protein-protein interactions with partner transcription factors not expressed in HEK 293 cells. In spite of this, the observed concordance of DMS activity scores with homeodomain structural models, the consistency of variant substitutions with the acidic exposure model of transcriptional activation domains, and the clinical variant classification performance all lend support to the validity of the DMS measurements.

## Materials and Methods

### Generation of the landing pad cell line

All cell lines were cultured in 90% DMEM (Gibco #11965092), 10% heat-inactivated fetal bovine serum (Gibco #16140089) supplemented with Penicillin-Streptomycin (Gibco #15140122). Unless otherwise noted, all transfections were carried out with cells plated at a density of 6×10⁵ cells/well in a six-well plate 24 hours prior to transfection. Each well of cells was transfected with the indicated plasmids and 4 µL each of Lipfectamine 3000 and P3000 reagent (Invitrogen #L3000015), suspended in 250 µL Opti-MEM (Gibco #31985062). For all transfections, the Lipofectamine 3000 was mixed with half the volume of Opti-MEM, while the P3000 reagent and DNA were combined in the remaining Opti-MEM. The Opti-MEM was mixed, and incubated at room temperature for 15 min, before being added dropwise to the prepared cells. All flow cytometry was performed on a Cyoflex S flow cytometer (Beckman-Coulter, V4-B2-Y4-R3 model).

In experiments where constructs were integrated into the landing pad, cells were transfected with a mixture of the indicated constructs and Addgene #11916, a plasmid expressing Cre recombinase. Three days post-transfection, media was replaced with fresh media supplemented with 5 nM AP1903/Rimiducid (MedChemExpress #HY-16046), which selects against the iCasp9 inducible caspase present in the naïve landing pad (Straathof et al. 2005). After 24 hours, media was replaced again, with fresh media supplemented with 5 nM AP1903 and 1 μg/mL puromycin (Sigma-Aldrich #P8833), which selects for the presence of puromycin N-acetyltransferase carried on the integrated construct. Surviving cells were allowed to recover and expand, with replating as needed to normalize cell density. Following integration, cells were continually grown in media supplemented with 5 nM AP1903 and 1 μg/mL puromycin.

To generate the landing pad cell line, HEK 293 cells were co-transfected with with 400 ng pJLS83, a plasmid carrying the landing pad sequence, and 400 ng Addgene #105927, a plasmid expressing Cas9 and an sgRNA targeting the human Rosa26 safe harbor locus. Eight days post-transfection, individual cells were sorted into single wells of multiple 96-well plates pre-filled with 150 µL of a 1:1 mixture of fresh media and 0.22 μm-filtered conditioned media exposed to wild-type HEK 293 cells in culture for 72 hours. The sort was performed on a Cytoflex SRT (Beckman-Coulter, V5-B2-Y5-R3 model); sorted cells were gated for positive blue channel (B450) fluorescence to select for expression of the mTagBFP2 fluorescent protein carried on the landing pad construct.

Over the course of three weeks, individual clones were allowed to expand and screened to identify lines with low-variance mTagBFP2 expression and normal growth characteristics and morphology. To screen out lines with integration of the landing pad construct on multiple alleles, clones were co-transfected with 600 ng Addgene #11916, a plasmid expressing Cre recombinase, and a 1:1 mixture of 200 ng pJLS119 and 200 ng pJLS120, plasmids carrying the mEmerald and mCherry2 fluorescent proteins, respectively, along with the necessary recombinase sites for landing pad recombination. Cells were measured by flow cytometry after AP1903+puromycin purification of LP-integrated cells, and any clones yielding cells exhibiting both green and red fluorescence were discarded. The final validated clonal landing pad cell line was frozen at low passage numbers, in CryoStor CS10 freezing medium (Biolife Solutions #210102).

### Generation of the CRX reporter cell line

A synthetic CRX reporter was synthesized, carrying eight repeats of a strong consensus CRX binding motif, CTAATCCC, each padded with a two base pair spacer motif (AG)(White et al. 2016). This reporter sequence was cloned upstream of the CMV-T6 minimal promoter(Loew et al. 2010) and the mEmerald fluorescent protein to produce pJLS96.

To produce lentivirus carrying the reporter, HEK 293T cells were seeded in a 10 cm dish, with 10 mL of media containing cells at a concentration of 3×10^5 cells/mL. Cells were transfected with mixture of 8 μg Addgene #12260, 1 μg Addgene #12259, 1 μg pJLS96, and 40 µL PEI in 500 µL Opti-MEM. 72 hours post-transfection, media was aspirated and concentrated using Lenti-X Concentrator (TaKaRa #631231) following the manufacturer’s recommended protocol. To create a “dead” reporter control, a second batch of lentivirus was produced using pJLS97, a variant of pJLS96 in which the CRX motifs were replaced with a variant known to abrogate CRX binding (CTACTCCC)(White et al. 2016).

Landing pad cells were separately transduced with the pJLS96 and pJLS97 lentiviruses. After 72 hours, the bulk populations of transduced cells were transfected with 400 ng pJLS38, a plasmid expressing human CRX. Three days post-transfection, individual pJLS96-transduced cells were sorted into single wells of multiple 96-well plates pre-filled with 150 µL of a 1:1 mixture of fresh medium and 0.22 μm-filtered conditioned media exposed to wild-type HEK 293 cells in culture for 72 hours. The sort was performed on a Cytoflex SRT (Beckman-Coulter, V5-B2-Y5-R3 model); sorted cells were gated for positive blue channel (B450) fluorescence to select for presence of the landing pad construct and for positive green fluorescence, relative to pJLS38-transfected cells transduced with the pJLS97 “dead” reporter construct.

Over the course of three weeks, individual clones were allowed to expand and screened to identify lines with low-variance mTagBFP2 expression and normal growth characteristics and morphology. Selected lines were co-transfected with 400 ng Addgene #11916 and 400 ng of either pJLS99, pJLS100, or pJLS101, plasmids carrying wild-type CRX, the p.R90W hypomorphic CRX variant, or miRFP670 (a transcriptionally inactive control protein), respectively. Following successful isolation of integrated cells, cells were screened by flow cytometry to maximize the dynamic range of mEmerald fluorescence between the pJLS99- and pJLS100-transfected cells. The final validated clonal landing pad + reporter cell line was frozen at low passage numbers, in CryoStor CS10 freezing medium.

### Cloning the CRX variant library

A DNA library comprising all possible human CRX single residue substitution variants was ordered as a combinatorial variant library from Twist Bioscience. The library construct was ligated between the AflII (NEB #R0520) and NheI (NEB #R3131) sites in pJLS84v2, a plasmid carrying recombinase sites matching the landing pad. The library fragment and plasmid backbone were mixed at a 3:1 molar ratio, targeting a total DNA amount of approx. 100 ng. The ligation was performed with T4 Ligase (NEB #M0202) at 16°C for 16 hours.

The ligation product was purified using a Monarch PCR & DNA Cleanup Kit (NEB #T1030) following the manufacturer’s recommended protocol, with elution in 6 μL H2O. The purified ligation product was transformed into 10-beta Electrocompetent E. coli (NEB #C3020). Briefly, 50 μL of electrocompetent cells were mixed with 5 µL purified ligation product on ice, and then split into two pre-chilled electroporation cuvettes (0.1 cm gap, Bio-Rad #1652089). Transformations were performed using a Bio-Rad GenePulser Xcell Electroporation System with PC Module (Bio-Rad #1652662), with the following conditions: 2000 V, 200 Ω, and 25 μF.

Following electroporation, 975 μL of pre-warmed 10-beta/Stable Outgrowth Medium was added (NEB #B9035) to each cuvette, and the outgrowths were pooled and transferred to a 14 mL round-bottom culture tube (Falcon #352059) for incubation with agitation at 37°C for 1 hour. 1:1000 and 1:10,000 dilutions of the outgrowth were plated on LB plates carrying 50 μg/mL kanamycin (LB+Kan50, Sigma-Aldrich #K1377) and incubated at 37°C for 16 hours. 1700 μL of the outgrowth was inoculated into 200 mL LB+Kan50 and incubated, with agitation, at 37°C for approx. 10 hours, until an OD600 of 3.0 was reached. Based on colony counts from the dilution plates, the 200 mL culture was inoculated with approx. 5.4 million colony-equivalents.

The 200 mL culture was centrifuged at 3100 ×g for 10 min to pellet the E. coli, and plasmid DNA was purified from the resulting cell pellet using a GenElute HP Plasmid Maxiprep Kit (Sigma-Aldrich #NA0310) following the manufacturer’s recommended protocol with elution in 3 mL H2O. The purified plasmid library was concentrated using an Eppendorf Vacufuge plus, yielding the “step one” CRX DMS library.

To add barcodes to this library, a short random barcoding oligo (“variant barcode”, “vBC”) was ordered as an Ultramer DNA Oligo from Integrated DNA Technologies (pJLS84v2+BC1). 1 μg of the step one plasmid library was linearized with XhoI (NEB #R0146) at 37°C for 16 hours. The digest product was run on a 1% agarose gel and the band was excised and purified using a Monarch DNA Gel Extraction Kit (NEB #T1020) following the manufacturer’s recommended protocol. The linearized plasmid library and barcoding oligo were assembled with NEBuilder HiFi DNA Assembly Master Mix (NEB #E2621) at a 1:5 molar ratio, targeting approx. 200 fmol total DNA in the assembly reaction. The assembly reaction was incubated at 50°C for 1 hour, and then purified using a Monarch PCR & DNA Cleanup Kit.

The purified, assembled, barcoded library was transformed following the same protocol as the step one DMS library. The 200 mL culture was inoculated with 40 μL of outgrowth and grown to an OD600 of 2.0 (approx. 13 hours). Based on colony counts from the dilution plates, the 200 mL culture was inoculated with approx. 85,000 colony-equivalents. Plasmid DNA was purified and concentrated using the same procedure as for the step one DMS library, yielding the “step two” CRX DMS library.

To add the puromycin N-acetyltransferase (PAC) cassette to the step two library, a fragment carrying an internal ribosome entry site (IRES) and PAC coding sequence was amplified from pJLS98 using primers JClPr88 and JClPr89. 1 μg of the step two plasmid library was linearized with SpeI-HF (NEB #R3133) at 37°C for 16 hours, and purified following the same protocol as the step two library. The IRES-PAC cassette and linearized step two library were assembled and purified following the same protocol used to generate the step two library.

The purified, assembled, PAC-containing library was transformed following the same protocol used to generate the step one DMS library. The 200 mL culture was inoculated with 1850 μL of outgrowth and grown to an OD600 of 3.5 (approx. 11 hours). Plasmid DNA was purified and concentrated using the same procedure as for the step one DMS library, yielding the “step three” CRX DMS library.

A second short random barcoding oligo (“random barcode”, “rBC”) was ordered as an Ultramer DNA Oligo from Integrated DNA Technologies (pJLS84v2+BC2). 1 μg of the step three plasmid library was linearized with XhoI at 37°C for 16 hours, and purified following the same protocol as the step two library. The barcoding oligo and linearized step three library were assembled and purified following the same protocol used to generate the step two library.

The purified, assembled, barcoded library was transformed following the same protocol used to generate the step one DMS library. The 200 mL culture was inoculated with 1850 μL of outgrowth and grown to an OD600 of 3.0 (approx. 10 hours). Plasmid DNA was purified and concentrated using the same procedure as for the step one DMS library, yielding the final CRX DMS library.

### Associating variants with barcodes

To associate CRX variants with barcodes, the step three CRX DMS library was sequenced using a PacBio Revio long-read sequencer. Briefly, 4 μg of the barcoded plasmid library was linearized by digestion with NruI-HF (NEB #R3192) at 37°C for 16 hours. The linearized library was purified using a Monarch PCR & DNA Cleanup Kit following the manufacturer’s recommended protocol with elution in 20 µL H2O. The purified linearized library was used to prepare a PacBio sequencing library using a SMRTbell prep kit 3.0 (PacBio #102-141-700), following the manufacturer’s recommended protocol (PacBio Protocol #102-166-600 REV02) with the following modifications: DNA shearing was skipped and the 20 μL of purified linearized library was used directly as input for Repair and A-tailing after the addition of 27 μL of Low TE buffer; the final cleanup with SMRTbell cleanup beads was not performed (no size selection). The prepared library was sequenced on a single Revio SMRT cell.

Reads were aligned to a synthetic reference sequence comprising the expected library plasmid structure with wild-type CRX and “N” nucleotides in place of the expected barcode location using minimap2 (v2.24, with parameters -A2 -B4 -O12 -E2 –end-bonus=13 –secondary=no –cs=long). The resulting PAF file was parsed with a custom Python script to generate a barcode-to-variant map. From 9.5 million total HiFi reads, 63.8% contained a valid barcode and a single missense CRX variant or wildtype sequence. The remaining reads represent a mixture of sequencing errors, low-quality reads, or valid reads of constructs that failed barcoding, acquired indels during cloning, or contained more than a single CRX missense variant. Of 75,946 observed barcodes on full-length plasmid reads, 86.8% mapped to a single CRX variant, while 7.3% mapped to wildtype CRX. Of the remaining barcodes, 4.9% mapped to CRX constructs with more than a single missense variant, while only approx. 1% of barcodes could not be unambiguously assigned—i.e. were observed to co-occur with different variants in different reads. All barcodes not uniquely mapping to a single CRX missense variant or wild-type CRX were discarded.

### Measuring variant activity

To conduct the DMS, landing pad + reporter cells were plated at a density of 3 × 10^5 cells/mL in 150 mm dishes, 30 mL per dish, three dishes per biological replicate. 24 hours later, each dish was transfected with 2 μg of the final CRX DMS library, as well as 8 μg Addgene #11916, 50 μL Lipofectamine 3000, and 50 μL P3000, in 3 mL Opti-MEM. Three days later, media was replaced with fresh media supplemented with 5 nM AP1903. Six hours later, media was replaced again with fresh media supplemented with 5 nM AP1903 and 1 μg/mL puromycin. Four days later, media was removed, Accutase (Innovative Cell Technologies #AT104) was added to dissociate cells, and cell suspensions from each of the three dishes for each biological replicate were pooled. Cell suspensions were centrifuged for 5 min at 150 ×g to pellet cells. Pelleted cells were resuspended in fresh media supplemented with 5 nM AP1903 and 1 μg/mL puromycin, and plated in a T-150 flask, one per replicate.

Cells were sorted into four bins based on reporter fluorescence on a Sony SY3200 fluorescence-activated cell sorter, recovering between 500,000 and 3,000,000 cells per bin. Sort distributions and gating strategies for each replicate are shown in Supplementary Source Data. Sorted fractions were pelleted by centrifugation for 5 min at 150 ×g, resuspended in 12 mL fresh media supplemented with 5 nM AP1903 and 1 μg/mL puromycin, and each plated in a fresh T-75 flask. 3–6 days post-sort, cell fractions were harvested by dissociation with 5 mL Accutase, centrifuged, and frozen in 1.5 mL microcentrifuge tubes.

Genomic DNA was extracted from each fraction using a Monarch Genomic DNA Purification Kit (NEB #T3010). Barcode sequences were amplified from gDNA using a two-step protocol: gDNA was first amplified with primers JLSPr141–144 and JLSPr165–168+171+172 using Q5 High-Fidelity DNA Polymerase (NEB #M0491) for 20 cycles with an annealing temperature of 65°C, and then with primers IDT10_i7_NN and IDT10_i5_NN, where NN is replaced with the unique indexing barcode ID, for sample multiplexing (10 cycles, 65°C annealing temperature). Prepared fraction amplicon libraries were sequenced on an Illumina NovaSeq X Plus instrument in a series of shared 10B flow cells, targeting 100 million reads per replicate.

Reads were cleaned with fastp (v0.23, with default parameters). Barcodes were extracted and counted using a custom script. vBCs were mapped to variants using the results of the PacBio long-read sequencing of the barcoded plasmid library. vBCs failing to match an expected barcode from the PacBio sequencing were error-corrected up to a Hamming distance of 1, if and only if they could be unambiguously mapped to an expected variant. All reads with non-mapping or ambiguous vBCs were discarded. vBC-rBC pairs with fewer than three reads were discarded. vBCs with fewer than 10 unique rBCs in any of the four bins were discarded. Raw activity scores were computed by summing the number of rBCs per vBC in each of the four bins divided by the sum of the number of rBCs across all four bins, weighted by the mean fluorescence intensity of each bin. Raw activity scores for each variant were divided by the wild-type CRX activity score, yielding the normalized DMS activity score used throughout this manuscript.

### Western blotting for CRX abundance

Cells were harvested by removal of media and addition of RIPA buffer (Cell Signaling Technology #9806, diluted to 1X in deionized water) supplemented with 1X Halt Protease Inhibitor Cocktail (Thermo Scientific #78430) and incubated on ice for 30 minutes. Cell lysate was centrifuged at 21,000 x g for 10 minutes and supernatant was mixed with Blue Loading Buffer (Cell Signaling Technology #7722) per the manufacturer’s recommended ratio. The combined supernatant and loading buffer was boiled for 8 min at 97°C and cooled to room temperature. Samples were run on 4–20% Mini-PROTEAN TGX Precast Protein Gels (Bio-Rad #4561096) in Tris/Glycine/SDS buffer (Bio-Rad #1610732, diluted to 1X in deionized water). Transfers were performed using a Bio-Rad Trans-Blot Turbo instrument, RTA Mini 0.2 µm PVDF Transfer Kit (Bio-Rad #1704272), and Trans-Blot Turbo Transfer Buffer (Bio-Rad #10026938). CRX (A-9) (Santa Cruz Biotechnology #sc-377138) and beta-Tubulin Rabbit Ab (Cell Signaling Technology #2146S) were used, along with Anti-rabbit IgG, HRP-linked (Cell Signaling Technology #7074S) and Anti-mouse IgG, HRP-linked (Cell Signaling Technology #7076S). TBST (EZ BioResearch #S-1012, diluted to 1X in deionized water) and EveryBlot Blocking Buffer (Bio-Rad #12010020) were used for washing and blocking, respectively.

### Analysis of transcriptional effector domain residue composition

The composition of transcriptional effector domains from CRX orthologs was compared to that of disordered regions found in annotated human TFs. CRX ortholog sequences were obtained from the UniProt UniRef50 cluster containing human CRX. Activating and bi-functional human TFs were curated by Soto et al. (2022). Disordered regions were predicted from these sequences using metapredict v2.6 (Emenecker et al. 2021). We compared the the fraction of D, E and F, W, Y, L residues in the predicted C-terminal IDRs of the CRX orthologs to the distribution observed for all IDRs predicted in the set of human TFs, evaluating a difference in the mean fraction by Mann-Whitney U Test.

### Local sequence effects of hydrophobic substitutions

Activity scores for substitutions occurring at non-hydrophobic residue positions in the transcriptional effector domain were fit to an interaction model between the class of substitution (hydrophobic or non-hydrophobic) and the number of immediately adjacent hydrophobic residues (0, 1, or 2). The following model was fit with ordinary least squares using the Python package statsmodels v0.14.0.

Normalized Activity ∼ (Class of Substitution) + (Number of Neighboring Hydrophobic Residues) + (Class of Substitution):(Number of Neighboring Hydrophobic Residues)

To determine the significance of the interaction term, a likelihood ratio test was performed against a reduced model excluding the interaction.

## Acknowledgements

We are grateful to Jessica Hoisington-Lopez and Maria-Lynn Crosby at the Washington University Center for Genome Sciences’ DNA Sequencing Innovation Lab as well as the staff at the Washington University Genome Technology Access Center for their assistance with the generation of high-throughput sequencing datasets. We are also grateful to the staff of the Alvin J. Siteman Cancer Center Flow Cytometry Core for their assistance with fluorescence-activated cell sorting. This work was supported by the National Institutes of Health: R01GM092910 and R21HG012146 to B.A.C., 1DP2CA290639 to A.S.H, and F30EY033640 to J.L.S.

## Data and Code Availability

Source code used for the analyses presented in this work is available at https://github.com/barakcohenlab/crx-dms-manuscript. Raw and processed sequencing data is available in GEO (GSE262060).

## Author Contributions

Investigation: J.L.S., D.M.G., J.L, Z.S., S.P.P; Writing — Original Draft: J.L.S.; Writing — Review & Editing: J.L.S., D.M.G., J.L., Z.S., S.P.P., A.S.H., M.A.W., B.A.C.; Supervision: A.S.H., M.A.W., B.A.C.

## Supplementary Figures

**Figure S1:**
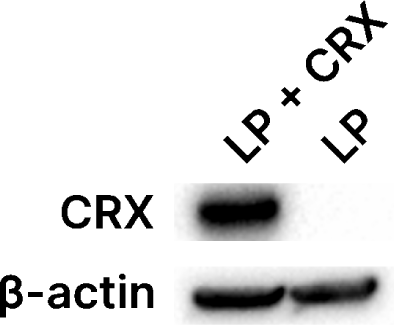
CRX expression in Landing Pad + Reporter cells. Western blot for CRX in HEK 293-derived LP + Reporter cells, either with wild-type CRX integrated in the landing pad (“LP + CRX”) or no CRX integrated, demonstrating a lack of expression of the endogenous copy of CRX.

**Figure S2:**
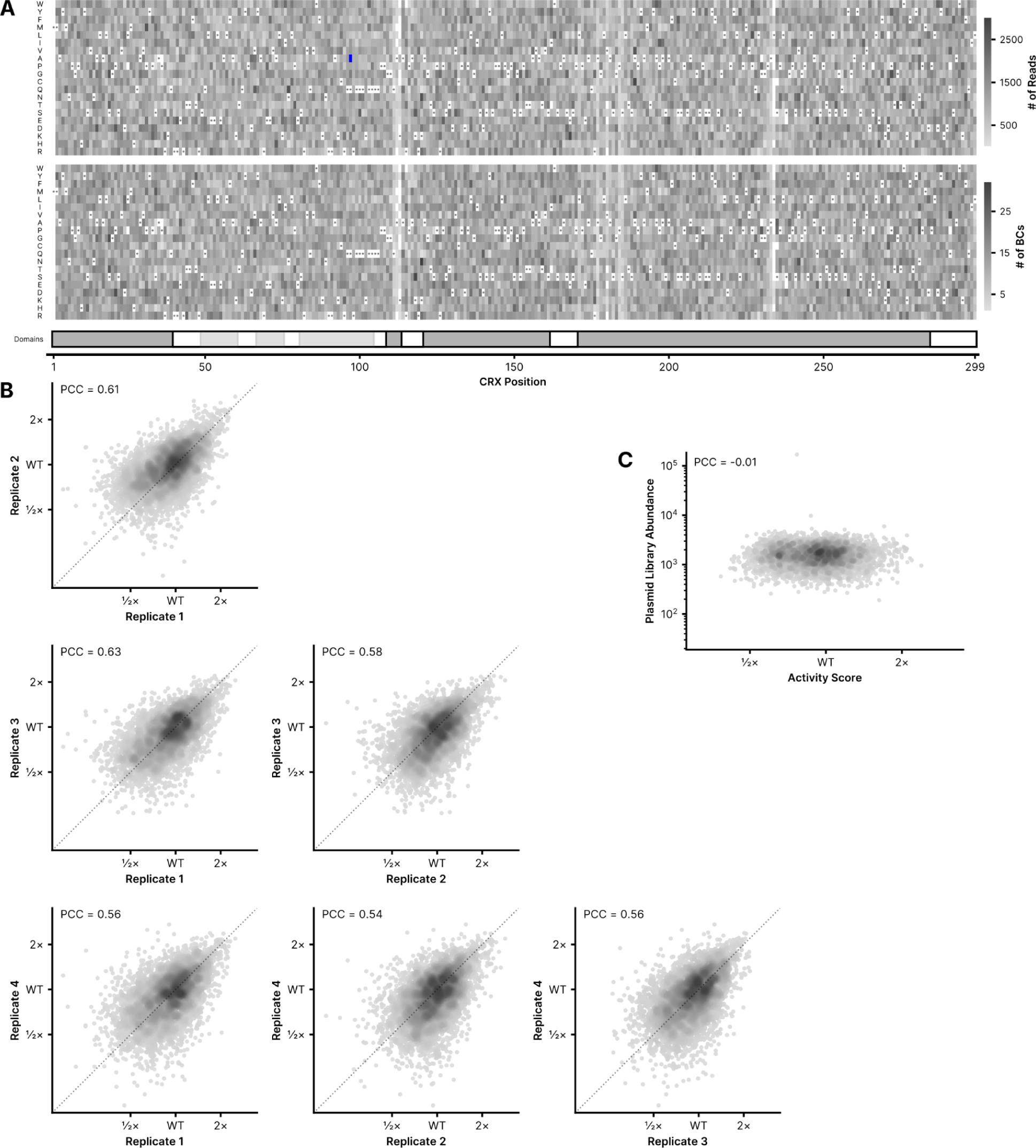
Library construction and quality control. (**A**) Number of reads (top) and barcodes (bottom) per variant as determined by PacBio long read sequencing of the cloned *CRX* variant library. The blue box indicates a single variant (Q97A; 10,305 reads) which was overrepresented in the library cloning process and is clipped in the colormap. Wildtype CRX was measured with 426,393 reads and 5,514 barcodes. (**B**) Variant activity score reproducibility across four replicates; DMS activity scores in the main text are a per-variant average of these four replicates. The dotted line in each plot is the line y=x, for visual reference. (**C**) Correlation between per-variant DMS activity scores and variant abundance in the input plasmid library.

**Figure S3:**
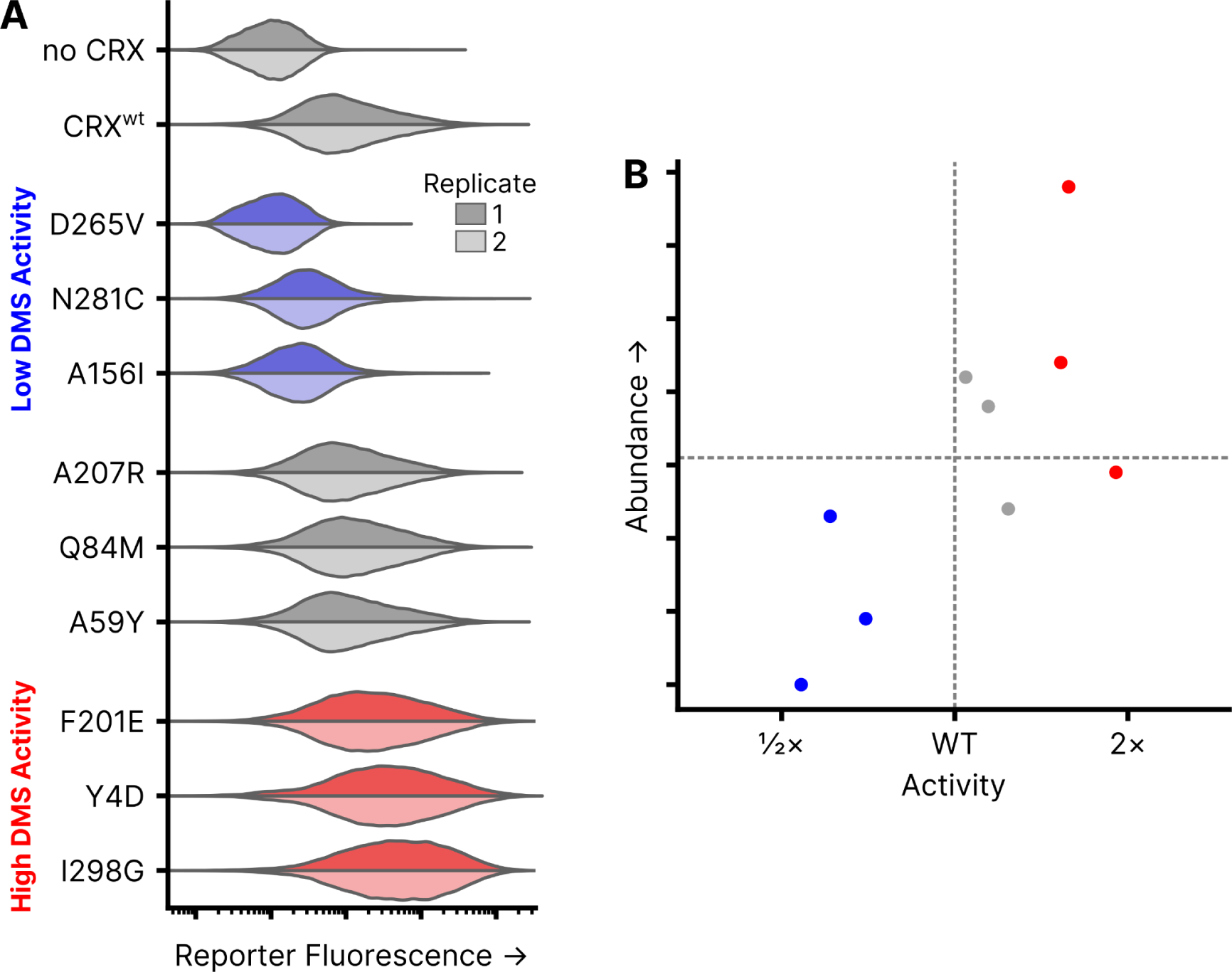
Per-variant activity and abundance relative to wild-type CRX. (**A**) Fluorescent reporter measurements (arbitrary units) of individual CRX variants integrated into the LP+Reporter cell line, separated by their classification in the DMS assay. Two independent biological replicate experiments per sample (distributions plotted from 40,000 cells). (**B**) Protein abundance of the variants in (A) measured by western blot quantification relative to β-tubulin (arbitrary units).

**Figure S4:**
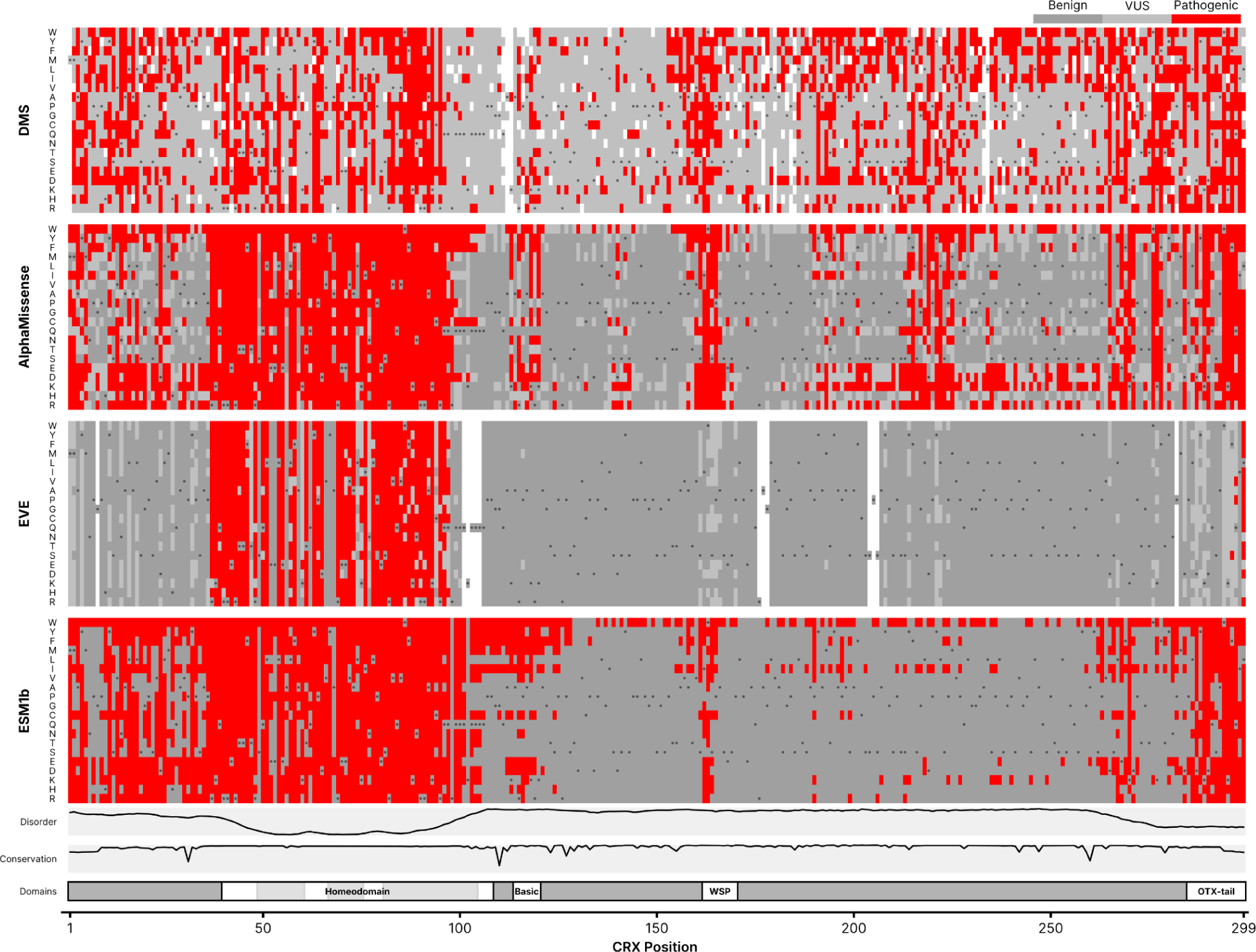
Comparison of CRX DMS results with computational variant effect predictors. For the DMS assay (top), any variant with an activity score significantly altered from wild-type CRX is called as “Pathogenic”, and other variants are called “VUS”. For AlphaMissense, model-produced classifications are shown. For EVE, the “75_pct_retained_ASM” classifications were used. For ESM1b, the recommended pathogenicity cutoff of -7.5 was used. Disorder, domains, and conservation are shown as in Figure 1.

