## Supplementary Source Data for "Mutational scanning of *CRX* classifies clinical variants and reveals biochemical properties of the transcriptional effector domain"

**Replicate 1**

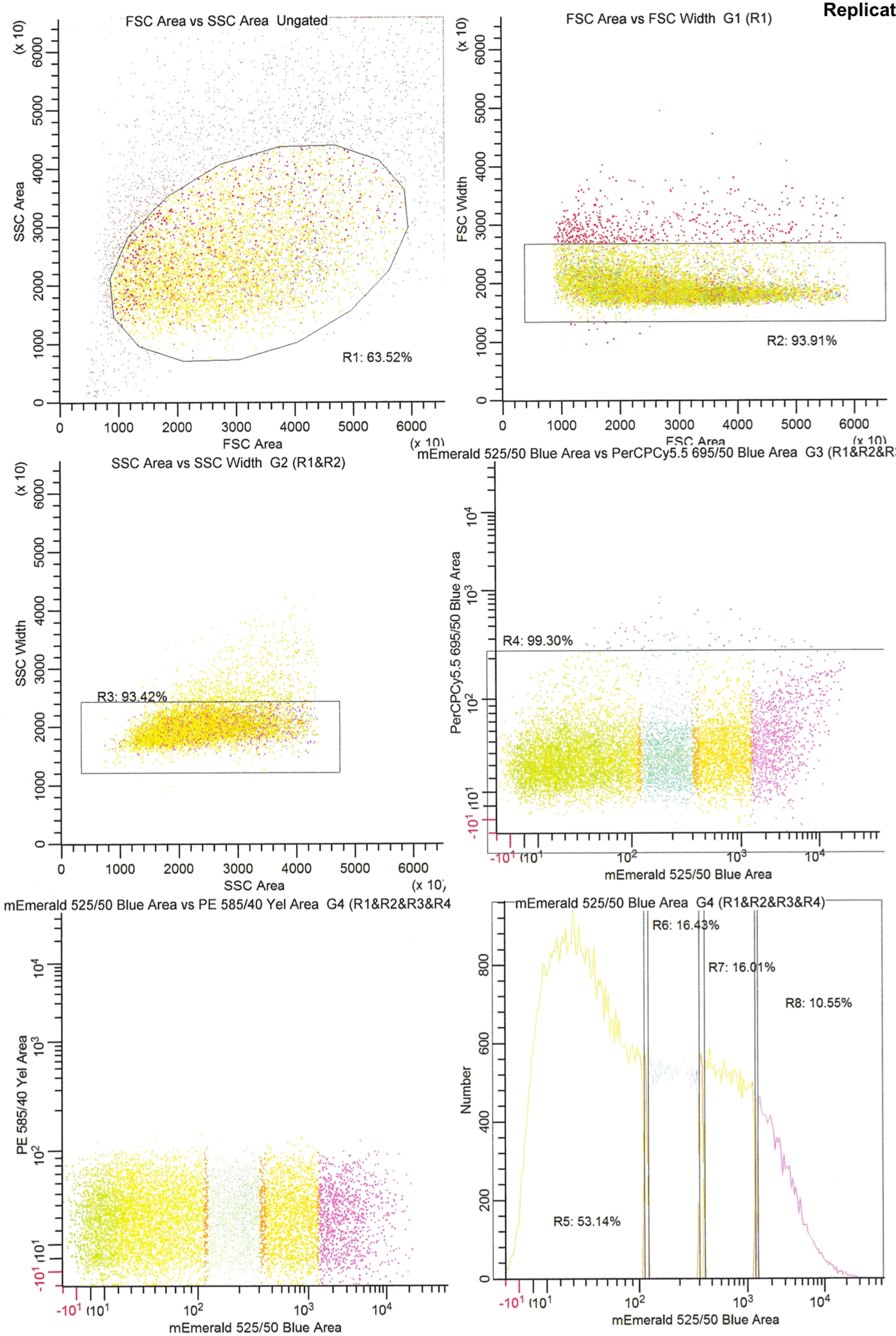

FSC Area vs SSC Area Ungated
SSC Area
(x 10)
FSC Area
(x 10)
R1: 63.52%
FSC Area vs FSC Width G1 (R1)
FSC Width
(x 10)
FSC Area
(x 10)
R2: 93.91%
SSC Area vs SSC Width G2 (R1&R2)
SSC Width
(x 10)
SSC Area
(x 10)
R3: 93.42%
mEmerald 525/50 Blue Area vs PerCPCy5.5 695/50 Blue Area G3 (R1&R2&R3)
PerCPCy5.5 695/50 Blue Area
mEmerald 525/50 Blue Area
R4: 99.30%
mEmerald 525/50 Blue Area vs PE 585/40 Yel Area G4 (R1&R2&R3&R4
PE 585/40 Yel Area
mEmerald 525/50 Blue Area
mEmerald 525/50 Blue Area G4 (R1&R2&R3&R4)
Number
mEmerald 525/50 Blue Area
R5: 53.14%
R6: 16.43%
R7: 16.01%
R8: 10.55%

**Replicate 1**

| Id | Percent | %Gate | geoMeanX | MedianX | LowX | HighX | geoMeanY |
|---|---|---|---|---|---|---|---|
| R5 | 29.40 | 53.14 | 37.07 | 36.12 | -29.90 | 108.81 | n/a |
| R6 | 9.09 | 16.43 | 179.85 | 180.54 | 116.09 | 301.65 | n/a |
| R7 | 8.86 | 16.01 | 624.75 | 623.24 | 341.70 | 1265.27 | n/a |
| R8 | 5.84 | 10.55 | 2836.04 | 2546.11 | 1357.29 | 63500.11 | n/a |

**Replicate 2**

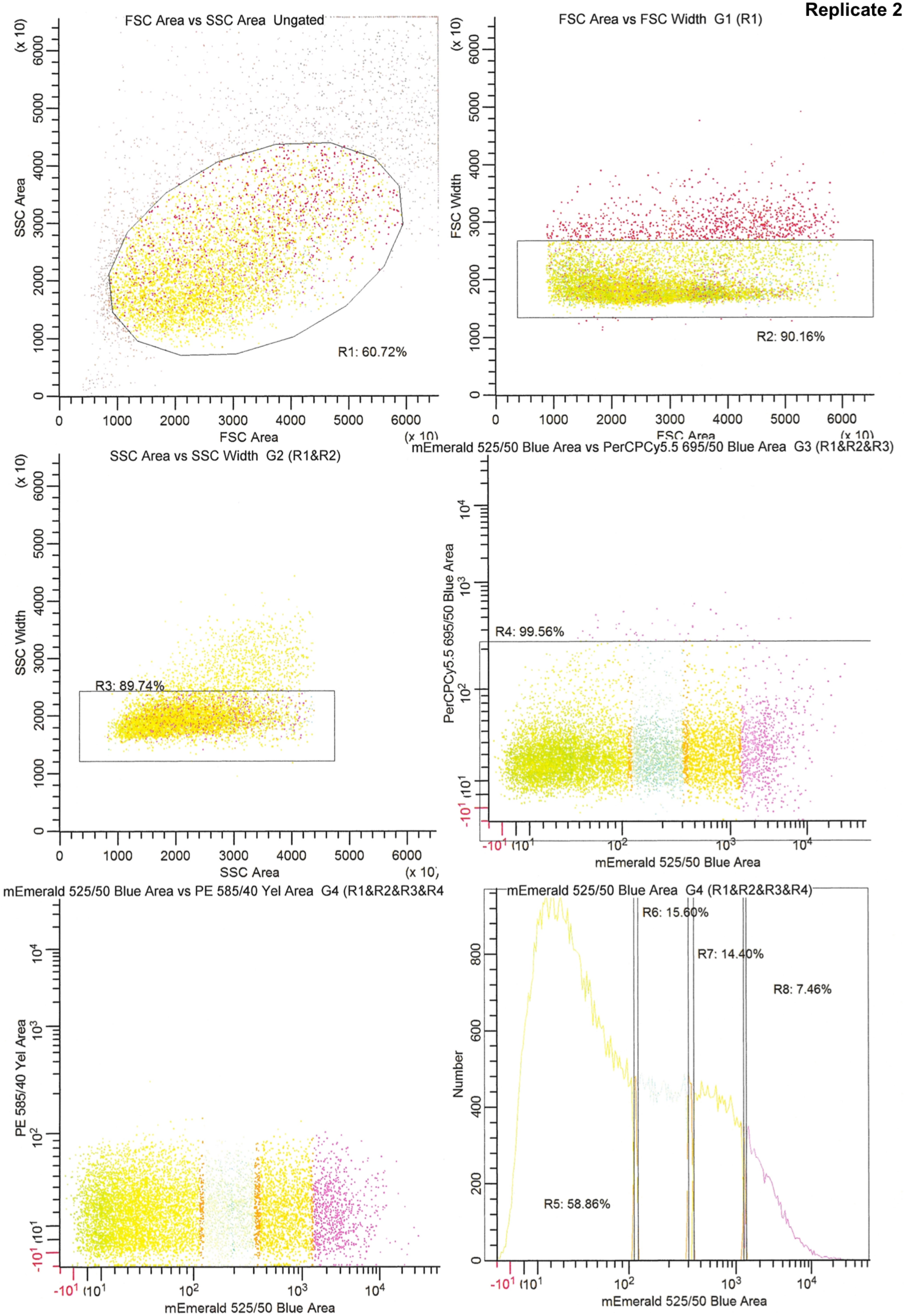

FSC Area vs SSC Area Ungated
SSC Area
FSC Area
(x 10)
R1: 60.72%
FSC Area vs FSC Width G1 (R1)
FSC Width
FSC Area
(x 10)
R2: 90.16%
SSC Area vs SSC Width G2 (R1&R2)
SSC Width
SSC Area
(x 10)
R3: 89.74%
mEmerald 525/50 Blue Area vs PerCPCy5.5 695/50 Blue Area G3 (R1&R2&R3)
PerCPCy5.5 695/50 Blue Area
mEmerald 525/50 Blue Area
R4: 99.56%
mEmerald 525/50 Blue Area vs PE 585/40 Yel Area G4 (R1&R2&R3&R4
PE 585/40 Yel Area
mEmerald 525/50 Blue Area
mEmerald 525/50 Blue Area G4 (R1&R2&R3&R4)
Number
mEmerald 525/50 Blue Area
R5: 58.86%
R6: 15.60%
R7: 14.40%
R8: 7.46%

**Replicate 2**

| Id | Percent | %Gate | geoMeanX | MedianX | LowX | HighX | geoMeanY |
|---|---|---|---|---|---|---|---|
| R5 | 28.79 | 58.86 | 35.72 | 33.71 | -29.90 | 108.81 | n/a |
| R6 | 7.63 | 15.60 | 179.40 | 179.86 | 116.09 | 301.65 | n/a |
| R7 | 7.04 | 14.40 | 619.24 | 614.29 | 341.70 | 1265.27 | n/a |
| R8 | 3.65 | 7.46 | 2659.13 | 2404.46 | 1357.29 | 63500.11 | n/a |

**Replicate 3**

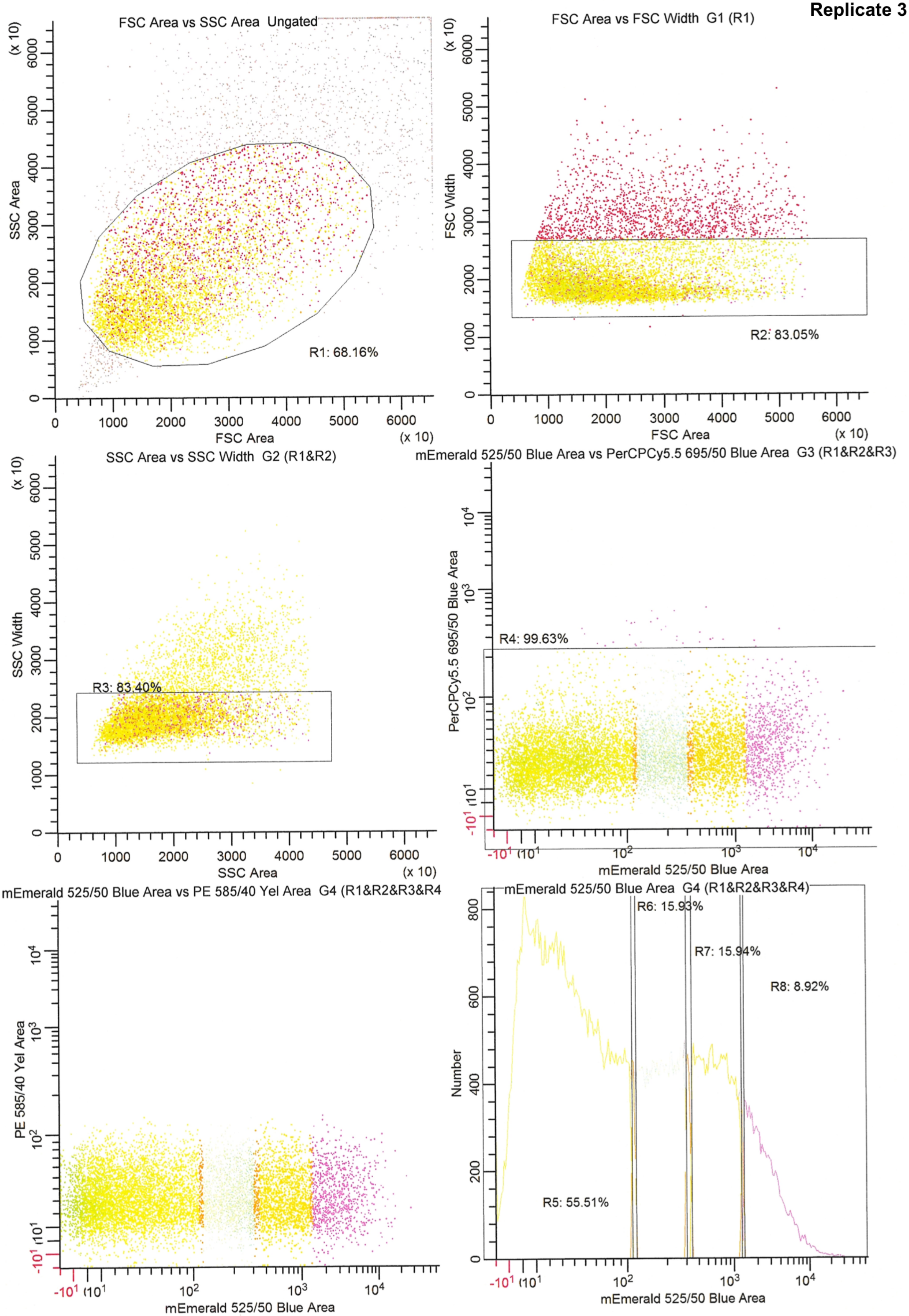
FSC Area vs SSC Area Ungated
SSC Area
(x 10)
FSC Area
(x 10)
R1: 68.16%
FSC Area vs FSC Width G1 (R1)
FSC Width
(x 10)
FSC Area
(x 10)
R2: 83.05%
SSC Area vs SSC Width G2 (R1&R2)
SSC Width
(x 10)
SSC Area
(x 10)
R3: 83.40%
mEmerald 525/50 Blue Area vs PerCPCy5.5 695/50 Blue Area G3 (R1&R2&R3)
PerCPCy5.5 695/50 Blue Area
mEmerald 525/50 Blue Area
R4: 99.63%
mEmerald 525/50 Blue Area vs PE 585/40 Yel Area G4 (R1&R2&R3&R4
PE 585/40 Yel Area
mEmerald 525/50 Blue Area
mEmerald 525/50 Blue Area G4 (R1&R2&R3&R4)
Number
mEmerald 525/50 Blue Area
R5: 55.51%
R6: 15.93%
R7: 15.94%
R8: 8.92%

**Replicate 3**

| Id | Percent | %Gate | geoMeanX | MedianX | LowX | HighX | geoMeanY |
|---|---|---|---|---|---|---|---|
| R5 | 26.10 | 55.51 | 31.46 | 29.39 | -29.90 | 108.81 | n/a |
| R6 | 7.49 | 15.93 | 181.46 | 182.74 | 116.09 | 301.65 | n/a |
| R7 | 7.50 | 15.94 | 623.93 | 623.07 | 341.70 | 1265.26 | n/a |
| R8 | 4.20 | 8.92 | 2723.14 | 2446.03 | 1357.29 | 63500.11 | n/a |

**Replicate 4**

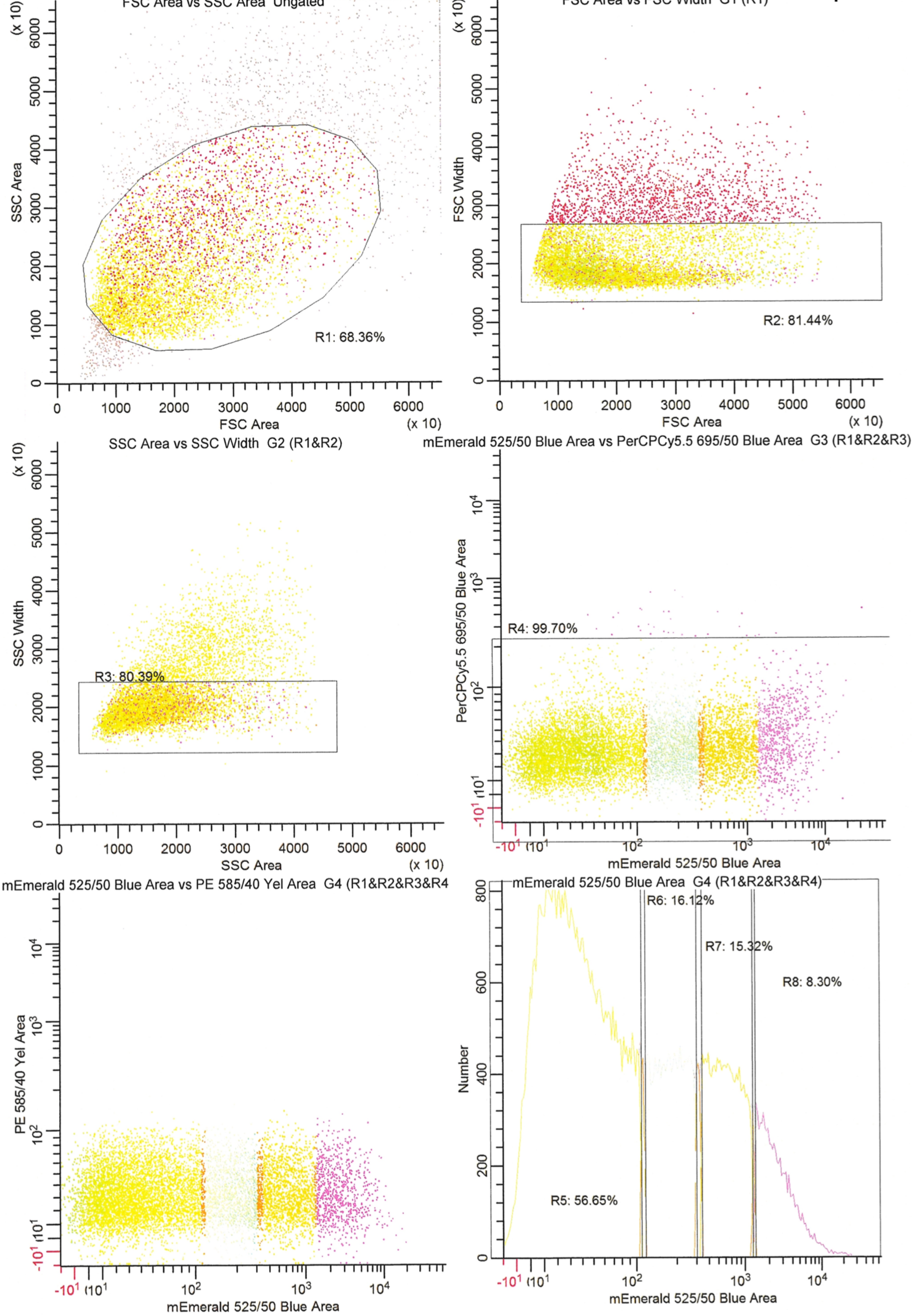
FSC Area vs SSC Area Ungated
SSC Area
FSC Area
(x 10)
R1: 68.36%
FSC Area vs FSC Width G1 (R1)
FSC Width
R2: 81.44%
SSC Area vs SSC Width G2 (R1&R2)
SSC Width
SSC Area
R3: 80.39%
mEmerald 525/50 Blue Area vs PerCPCy5.5 695/50 Blue Area G3 (R1&R2&R3)
PerCPCy5.5 695/50 Blue Area
R4: 99.70%
mEmerald 525/50 Blue Area
mEmerald 525/50 Blue Area vs PE 585/40 Yel Area G4 (R1&R2&R3&R4
PE 585/40 Yel Area
mEmerald 525/50 Blue Area G4 (R1&R2&R3&R4)
Number
R5: 56.65%
R6: 16.12%
R7: 15.32%
R8: 8.30%

**Replicate 4**

| Id | Percent | %Gate | geoMeanX | MedianX | LowX | HighX | geoMeanY |
|---|---|---|---|---|---|---|---|
| R5 | 25.28 | 56.65 | 34.76 | 32.71 | -29.90 | 108.81 | n/a |
| R6 | 7.19 | 16.12 | 180.59 | 181.62 | 116.09 | 301.65 | n/a |
| R7 | 6.83 | 15.32 | 619.20 | 616.98 | 341.70 | 1265.26 | n/a |
| R8 | 3.70 | 8.30 | 2674.46 | 2399.26 | 1357.29 | 63500.11 | n/a |
